# Single-cell copy number lineage tracing enabling gene discovery

**DOI:** 10.1101/2020.04.12.038281

**Authors:** Fang Wang, Qihan Wang, Vakul Mohanty, Shaoheng Liang, Jinzhuang Dou, Jincheng Han, Darlan Conterno Minussi, Ruli Gao, Li Ding, Nicholas Navin, Ken Chen

## Abstract

Aneuploidy plays critical roles in genome evolution.

Alleles, whose dosages affect the fitness of an ancestor, will have altered frequencies in the descendant populations upon perturbation.

Single-cell sequencing enables comprehensive genome-wide copy number profiling of thousands of cells at various evolutionary stage and lineage. That makes it possible to discover dosage effects invisible at tissue level, provided that the cell lineages can be accurately reconstructed.

Here, we present a Minimal Event Distance Aneuploidy Lineage Tree (MEDALT) algorithm that infers the evolution history of a cell population based on single-cell copy number (SCCN) profiles. We also present a statistical routine named lineage speciation analysis (LSA), which facilitates discovery of fitness-associated alterations and genes from SCCN lineage trees.

We assessed our approaches using a variety of single-cell datasets. Overall, MEDALT appeared more accurate than phylogenetics approaches in reconstructing copy number lineage. From the single-cell DNA-sequencing data of 20 triple-negative breast cancer patients, our approaches effectively prioritized genes that are essential for breast cancer cell fitness and are predictive of patient survival, including those implicating convergent evolution. Similar benefits were observed when applying our approaches on single-cell RNA sequencing data obtained from cancer patients.

The source code of our study is available at https://github.com/KChen-lab/MEDALT.

## Introduction

Aneuploidy, the phenomenon that genomes acquire or lose chromosomal fragments, has been causally implicated in a wide variety of human diseases such as neuropsychiatric disorders and cancer^1–3^. Genetic and phenotypic plasticity resulting from aneuploidy evolution causes treatment resistances and disease recurrences^4–6^, which fundamentally challenges current medicine. Recent studies have shown that not only disease tissues, but also pathologically normal tissues may contain a high degree of somatic mosaicisms (e.g., peripheral blood^7^ and esophagus^8^). Therefore, defining which copy number alterations (CNAs) cause pathogenesis and which are part of normal variations becomes increasingly important in genome medicine, especially for cancer^9, 10^.

Various efforts have been made to obtain comprehensive knowledge of CNAs responsible for cancer diagnostics, prognostics and targeted therapeutics. Systematic CNA analysis in over 10,000 primary tumor samples in the cancer genome atlas (TCGA) and 2,500 samples in the International Cancer Genome Consortium (ICGC) revealed distinct CNA landscapes in different cancer types ^11–13^. Comparison of CNAs amongst autologous tumors obtained at different stage from different histology revealed that CNAs are critical for tumor evolution across time and space. However, studies based on bulk tissue samples cannot fully depict the history of tumor evolution, which occurs in single-cell resolution^14^, and thus have limited power to discover the associated genetic drivers.

Recent advances in single-cell DNA sequencing (e.g., tagmentation based approach^15^ and single-cell CNV solution by the 10X Genomics) have enabled large-scale acquisition of single-cell copy number (SCCN) profiles in tens of thousands of cells at around 100 kb resolution (~0.1X sequencing coverage per cell)^16–19^. Other platforms such as single-cell RNA-sequencing^20, 21^ and single-cell ATAC-sequencing^22^ have also been utilized for SCCN profiling.

These SCCN profiles not only present a rich pool of genetic perturbations that are invisible at tissue level, but also potentiate reconstruction of cellular lineage, based on which the impact of an allele on cellular fitness can be measured. Thus, statistical approaches that integrate cellular lineage tracing with population genetic analysis^23^ can enable discovery of novel disease genes and mechanisms of disease progression.

So far, studies performing retrospective lineage tracing from single-cell data have largely been utilizing phylogenetics approaches designed to model species evolution, which is quite different from cellular evolution in terms of duration, scale, genetics and dynamics^24, 25^. Many existing phylogenetics approaches assume that genomic sites evolve independently and follow the so-called infinite site assumption (ISA)^26^. But in the context of aneuploidy, a genome site can often be altered repeatedly by different CNAs, due partly to constraints on genome and chromatin structures, properties of DNA replication/repairing^27^ and functional selection. To apply conventional maximum parsimony approaches on SCCN data, one has to over-segment genomic regions and represent copy numbers as characters in disjoint intervals, which ill-represents the properties of DNAs and distorts evolution propensity across copy number states. Other conventional methods using Euclidean, Hamming or correlational distances also ill-represent the segmental, non-linear nature of CNA evolution^28^, leading to inaccurate inference of tree topology and branch lengths. A few new phylogenetics approaches have been developed to tackle these limitations^29^. Particularly, the MEDICC^29^ algorithm infers a copy number phylogenetic tree from the allelic copy number profiles of a set of samples. However, the problem is NP-hard. Even the simplified solutions could be applied to only tens of genomes^30^ and are not scalable to current single-cell datasets consisting of thousands of cells.

## Results

### Overview of the methods

To address these challenges, we propose a new computational framework that performs lineage tracing from SCCN data and detects significant focal (gene resolution) and broad (chromosomal-arm resolution) CNAs associated with lineage expansion (**Fig. 1**).

**Fig. 1.**
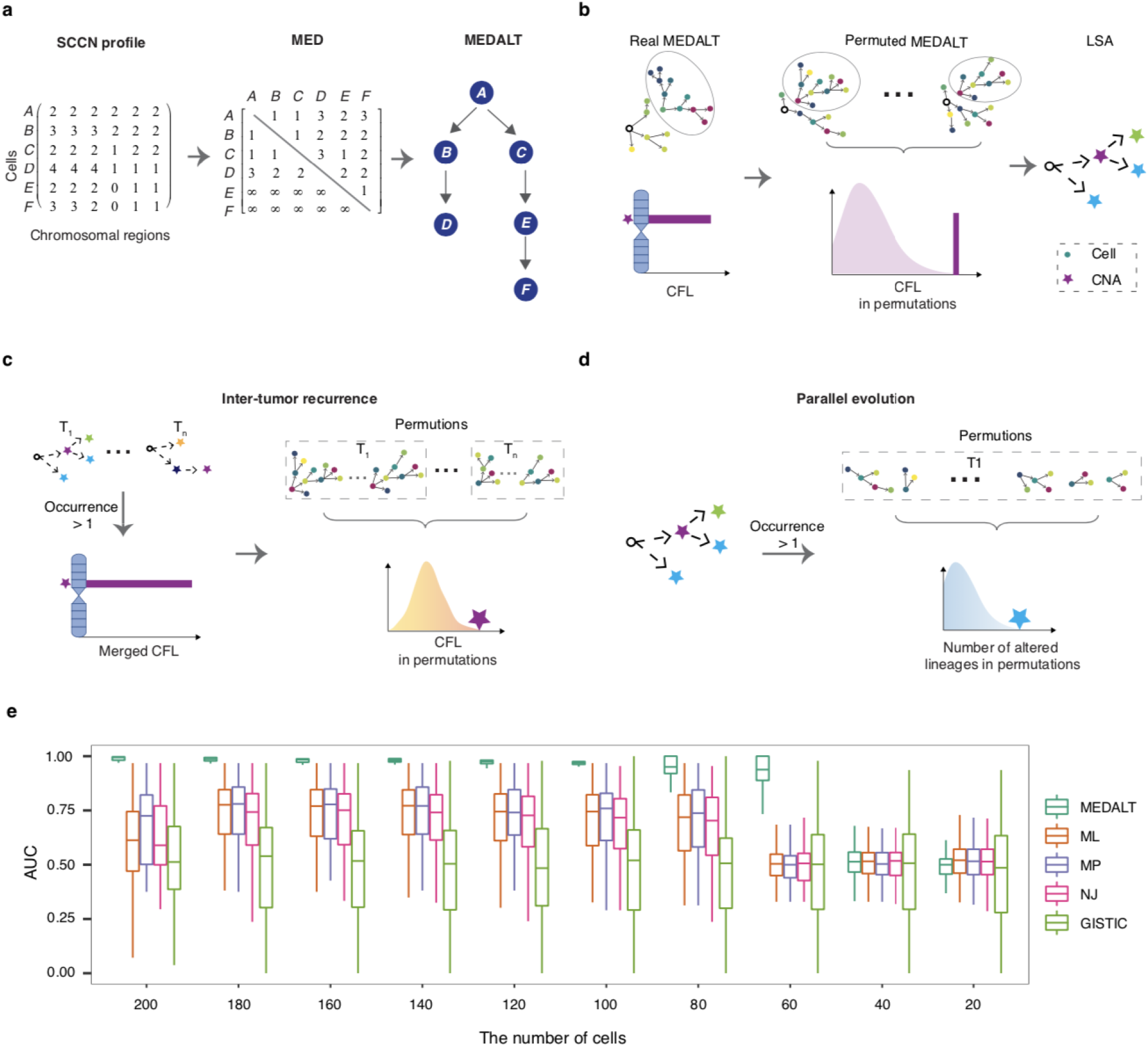
Algorithm flowchart and evaluation. **a.** Algorithm for constructing a MEDALT. **b.** Identification of non-random fitness-associated CNAs in an individual sample. **c.** Identifying non-random fitness-associated CNAs in a cohort of samples. **d.** Identifying parallel/convergent evolution CNAs in an individual sample. **e.** AUC of FAA identification based on 100 synthetic datasets. MP: maximal parsimony tree. NJ: neighbor-joining tree. ML: maximum likelihood tree.

We first collapse the given SCCN profiles into a data matrix, in which each row represents a cell and each column a chromosomal region. We then employ a distance metric called Minimal Event Distance (MED) to deduce the number and the series of single-copy gains or losses that are required to evolve the genome of one cell to the next (**Fig. S1a**) using an efficient greedy algorithm (**Methods** and **Table S1**).

We then infer a directed minimal spanning tree, named Minimal Event Distance Aneuploidy Lineage Tree (MEDALT, **Fig. 1a**), using an adapted version of the Edmond’s algorithm that scales polynomially with respect to the number of cells (**Methods** and **Table S2**). In a MEDALT, each node represents a cell, each edge represents a kinship between two cells, arrows point towards younger cells, and the root represents a normal diploid cell.

MEDALT allows a genomic region to be repetitively altered by multiple single-copy gains or losses. It provides a parsimonious interpretation, the minimal number of single-copy gains or losses that may have led to the evolution of the entire cell population.

An important constraint is that chromosomal fragments cannot be recovered if completely lost. To reflect that property, the MEDs originating from cells containing homozygous copy number loss are set to infinity.

Since MEDALT describes copy number evolution by segments instead of sites, we expect that it will enable more accurate cellular lineage tracing than do conventional phylogenetics methods (**Fig. S1b, Methods**).

We further establish a statistical routine, named Lineage Speciation Analysis (LSA), to prioritize CNAs and genes that are non-randomly associated with lineage expansion and thereby have potential functional impact.

To perform LSA, we first iteratively partition cells into lineages (subsets) based on the topology of the lineage tree. For each CNA region in each candidate lineage, we calculate a cumulative fold level (CFL) as the summation of the copy number levels in constituent cells (**Fig. S1c**). We then assess the statistical significance of the observed CFL with respect to a background distribution established from random lineages of similar sizes obtained from a permutation process (**Methods, Fig. 1b and Fig. S1d**). The permutation process randomly assigns SCCN profiles by chromosomes into different cells and reconstruct a lineage tree from each permuted dataset using the same lineage tracing algorithm. It is important to account for background variations induced by factors unrelated to cellular fitness such as high CNA prevalence at fragile sites or repeats that are non-functional, as shown in previous studies^31, 32^ and to account for bias of lineage tracing algorithms (**Supplementary Note**). The efficiency of MEDALT algorithm makes it possible to perform a large number of permutations in order to obtain a reasonably accurate background distribution. The statistically significant CNAs and genes so identified may not be causal themselves, but are associated with (e.g., co-occur) with causal fitness-impacting alterations. Thus, LSA distills the massive genome-wide SCCN data into a compact molecular blueprint, consisting of CNAs/genes occurring non-randomly at important moment during the course of the evolution with significant impact on the fitness of the descendant cells.

LSA can also be applied at cohort level to analyze single-cell data obtained from multiple patient samples. In that setting, the method creates meta-lineages combining cells from different patients and prioritizes events non-randomly occurring across background lineages established over the entire cohort (**Fig.1c** and **Methods**). Genes that are altered nonrandomly in multiple patients will likely have higher scores than those altered in a single patient.

Additionally, LSA can be applied to prioritize CNAs associated with parallel/convergent evolution^33^ (abbr. PLSA) by estimating the chance of a CNA occurring nonrandomly in two or more parallel lineages, as a consequence of positive selection (**Fig.1d** and **Methods**). This opens a new way for gene discovery that was substantially underpowered in bulk sample studies.

### *In silico* evaluation

To evaluate our approaches, we simulated copy number evolution in single cells using a Markov process parameterized by cell fitness parameters (**Fig. S2, Methods**)^34^. Spiked in randomly were fitness-associated alterations (FAAs), which indicate fitness change in a cell triggering subsequent lineage expansion. Synthetic SCCN profiles were created mimicking various CNA mechanisms such as genome doubling, breakage-fusion-bridge (BFB), tandem duplication, terminal deletion, unbalanced translocation, etc^27^. We created 100 simulated datasets, each containing around 200 cells. Besides obtaining MEDALTs, we also obtained phylogenetic trees using conventional maximum likelihood (ML), maximum parsimony (MP) and neighbor joining (NJ) approaches (**Methods**). In addition, we ran GISTIC^31^ (**Methods**), a method developed to prioritize CNAs in tissue samples by treating the cells as unrelated samples.

We then perform FAA detection in each dataset by performing LSA on individual trees inferred by various methods. We compare the detection performances using area under receiver operating characteristic curves (AUC, **Methods**). Overall, the MEDALT approach achieved substantially better detection performance than the other methods **(Fig. 1e)**. The benefits appeared robust over a range of cell numbers, when we repeated the benchmarking on subsets of the cells via random down-sampling, until the number of cells dropped below 60 (**Fig. 1e**).

### Detecting fitness-associated CNAs in disease cohorts

We applied our methods on the single-cell DNA-sequencing data acquired from 20 triple-negative breast cancer patients (TNBCs, **Table S3**)^16, 18^. SCCN profiles were generated using a variable binning method followed by circular binary segmentation (CBS)^35^ (**Fig. S3, Methods**). We obtained both MEDALTs and phylogenetic trees for each sample and ran LSA to identify non-random alterations at both sample and cohort levels.

We then compared the accuracy of the trees in inferring cellular timing using data from 4 patients with longitudinal pre-, mid- and post-treatment (neoadjuvant chemotherapy) samples. We found that MEDALTs ordered cells much more consistently with their biopsy timing than did the phylogenetic trees (**Fig. S4**), with pre-treatment cells appearing near the root and post-treatment cells near the leaves.

Consistent with previously studies^16, 18^, most of the TNBC samples appeared to have developed through branched evolution via multiple parallel lineages. Interestingly, the MEDALTs indicated that these parallel lineages may have distinct mutation rates **(Fig. 2a and b, Fig. S5)**, which may be attributable to variable degree of DNA damage repair (DDR) loss (**Fig. 2b, Methods**)^36^. Indeed, when we performed gene set enrichment analysis on genes identified by LSA, we found that the lineages of higher CNA rates have more DDR genes affected by the CNAs than the lineages of lower CNA rates **(Fig. S6**).

**Fig. 2.**
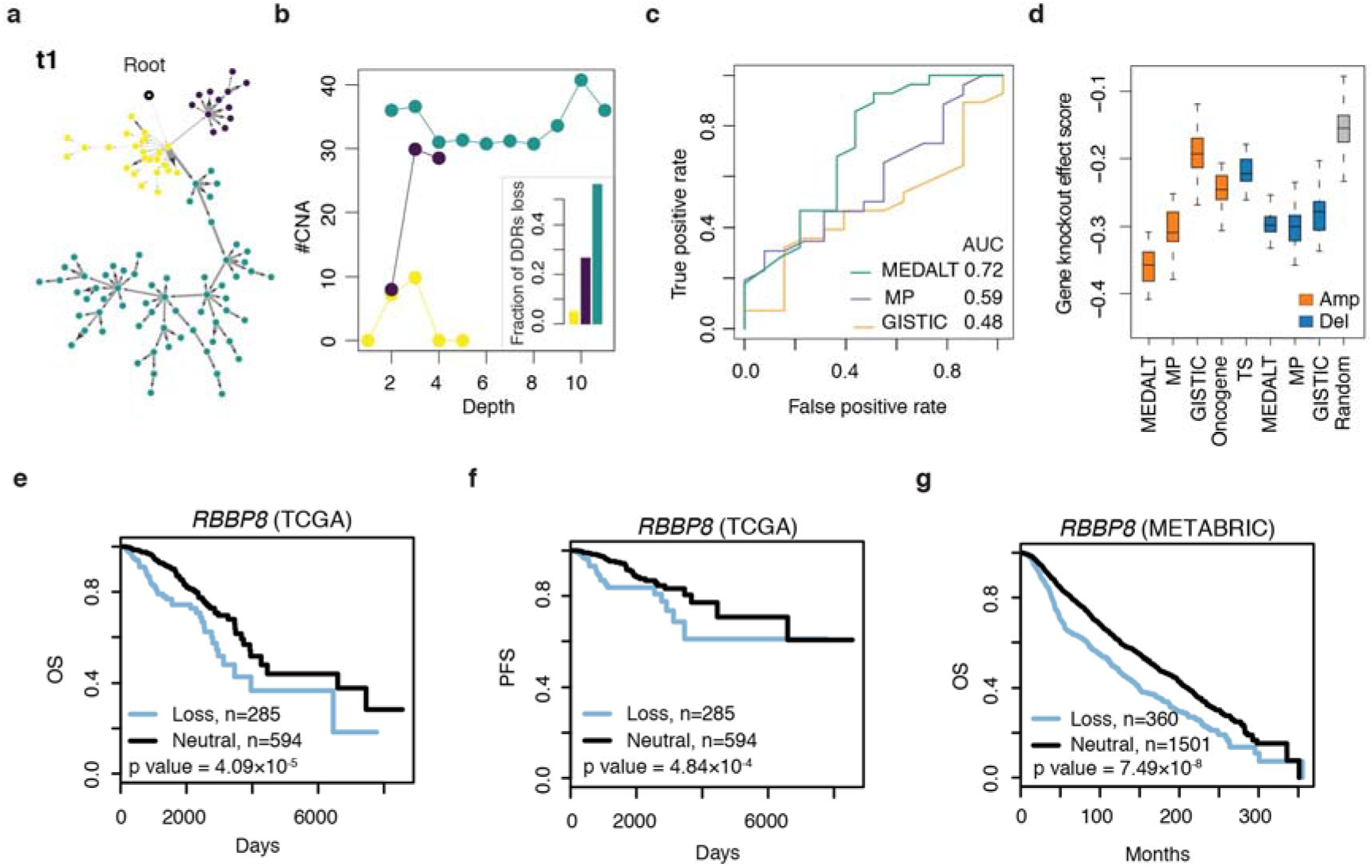
Application on scDNA-seq data from TNBCs. **a.** The MEDALT Inferred from patient t1. The widths of the edges are drawn proportional to the MEDs. Colors (yellow, blue and green) highlight the 3 main branches. **b.** The relationship between CNA numbers (Y-axis) and the depth on the tree (or distance to the root, X-axis). The barplot shows the fraction of DDR genes among the genes with copy number losses in the 3 lineages. **c.** ROC curve for identifying functionally important, broad CNAs in the literature (**Table S4**). **d.** The gene knockout effect scores of the gene sets (cohort LSA test p value < 0.001) identified based on the MEDALTs, MP trees and GISTIC in 29 breast cancer cell lines. Included as controls are 100 sets of 197 randomly selected genes, 234 oncogenes and 269 tumor suppressors (TS) in oncoKB and intOGen. The overall survival (OS) (**e**) and the Progression free survival (PFS) (**f**) of RBBP8 loss in TCGA breast cancer patients. **g.** OS of *RBBP8* loss in the METABRIC.

We identified fitness-associated CNAs at chromosomal and gene resolution using cohort-level LSA (*p* value < 0.001). For benchmarking, we also performed the same LSA on the MP trees. We also ran GISTIC^31^ on the pseudo-bulk copy number profiles generated by averaging the SCCN profiles across the cells in each sample (**Methods**).

Overall, the MEDALT plus LSA approach identified 30 broad CNAs, 80% of which have been functionally associated with breast cancer development and treatment outcome in the literature (**Table S4**). The accuracy was at least 13% higher than the results derived by the other methods (**Fig. 2c, Methods**). At gene resolution, our approach identified 197 genes, including 109 amplified and 88 deleted genes (**Supplementary Data 1**). In contrast, the MP plus LSA approach identified 130 genes, 82 of which were amplified and 48 deleted. GISTIC identified 60 genes, 33 of which were amplified and 27 deleted.

By examining the CRISPR knockout screen data in 29 breast cancer cell lines in the DepMap database^37^, we found that the 109 amplified genes identified by the MEDALT plus LSA approach had significantly lower gene knockoff effect scores than those of the 82 amplified genes detected based on the MP trees (one-side Wilcoxon rank-sum test, *p* = 2.75 × 10^−9^) and of the 33 genes detected by GISTIC (one-side Wilcoxon rank-sum test, *p* = 6.65 × 10^−17^) (**Fig. 2d**). The scores were also significantly lower than those of oncogenes (one-side Wilcoxon rank-sum test, *p* = 1.12 × 10^−15^) and tumor suppressors (one-side Wilcoxon rank-sum test, *p* = 2.81 × 10^−16^) reported in the oncoKB^38^ and intOGen^39^ databases, which are not specific to TNBC, and sets of randomly selected genes of identical size (one-side Wilcoxon rank-sum test, *p* = 8.97 × 10^−21^). Not significant were the scoring differences amongst the sets of deleted genes, due likely to challenges in calling deletions from noisy low-coverage data and in quantifying deleterious effects in lineages of limited cell numbers.

Among the 197 genes MEDALT nominated, some are not reported in the oncoKB^38^, COSMIC^40^ and intOGen^39^ databases (**Supplementary Data 2**) but supported by functional genomics data in large-scale cancer patient studies (**Fig. S7a**). For example, loss of *RBBP8* indicated worse prognosis among the breast cancer patients in TCGA and those in the METABRIC^41^ (**Fig. 2e** to **g**). *RBBP8* is a potentially interesting target as it interacts with *BRCA1* and modulates its function in transcriptional regulation, DNA repair and/or cell cycle checkpoint control^42^. In addition, loss of *PPP4R1* indicated worse prognosis in TCGA and the METABRIC as well (**Fig. S7b** to **d**).

In addition, we identified 107 genes that were likely positively selected (PLSA *p* value < 0.001, **Supplementary Data 3**) by convergent evolution in 7 of the 20 patients (**Fig. 3a**), by performing PLSA on the MEDALTs derived from individua patients. Among these, 65 genes were amplified. By repeating the same PLSA on the MP trees, we identified 355 genes, 252 of which were amplified. The set of 65 genes identified from the MEDALTs had significantly lower gene knockout effect scores (thus more essential) than those of the set of 252 genes identified from the MP trees (one-side Wilcoxon rank-sum test, p value = 4.07 × 10^−9^), of known oncogenes (one-side Wilcoxon rank-sum test, *p* = 2.81 × 10^−16^) and sets of randomly selected genes (one-side Wilcoxon rank-sum test, *p* = 9.01 × 10^−21^), based on the CRISPR screens of the 29 breast cancer cell lines in the DepMap^37^ (**Fig. 3b**). No significant scoring differences were found between the deleted genes identified from the MEDALTs and those identified from the MP trees, although both sets appeared more essential than the sets of known tumor suppressors and randomly selected genes.

**Fig. 3.**
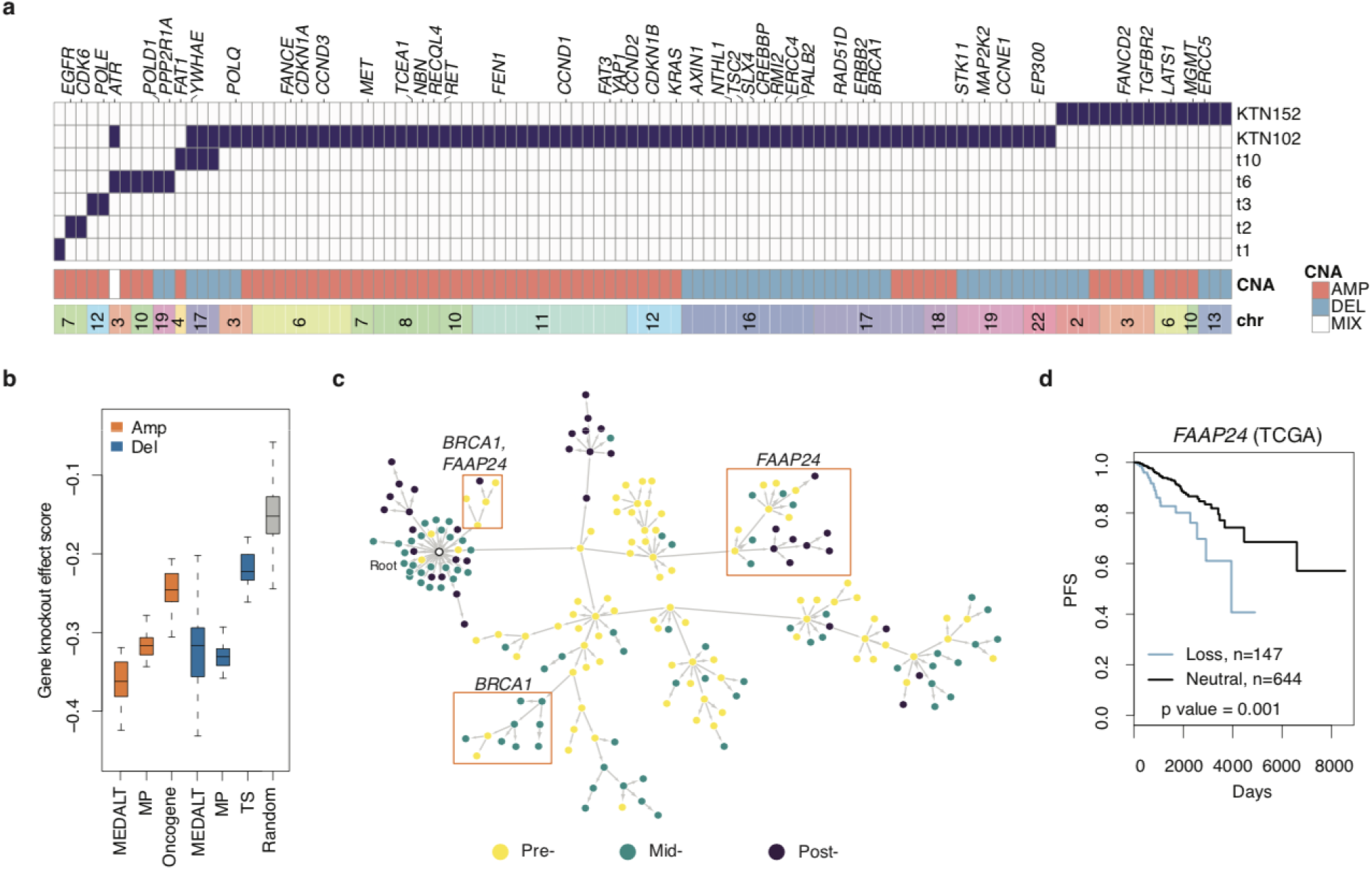
Convergent evolution in TNBCs. **a.** Genes associated with convergent evolution in the 20 TNBC patients. Labeled at the top are known cancer genes. AMP: copy number amplification, DEL: copy number deletions and MIX: genes with amplifications and deletions. **b.** The gene knockoff effect scores of gene set identified from the MEDALTs and the MP trees by PLSA (p value 0.001). Included as controls are 100 sets of 107 randomly selected genes, 234 oncogenes and 269 tumor suppressors (TS) in oncoKB and intOGen. **c.** The MEDALT of patient KTN102. Orange boxes highlight lineages under potential convergent evolution. **d.** Progression free survival of breast cancer patients with and without *FAAP24* loss in TCGA.

Among the 107 genes identified by PLSA, 42% were known cancer genes, a fraction higher than what we obtained from the cohort-level single-lineage LSA (38%, **Fig. S7e**). Loss of *FAAP24* appeared in two distinct lineages in patient KTN102 and was associated with worse progression free survival (PFS) in TCGA breast cancer data (**Fig. 3c** and **d**). Loss of *BRCA1* was also found in two parallel lineages, which were depleted of cells from the post-treatment sample (**Fig. 3c**). That observation may be explained by the fact that BRCAness tumors often respond to neoadjuvant chemotherapy^43, 44^.

### Applications on single cell RNA sequencing data

Our approaches are likely beneficial to characterizing SCCN data derived from single-cell RNA sequencing (scRNA-seq) experiments. To examine that possibility, we collected data obtained from paired primary and metastasis (or relapse) samples of a variety of cancer patients, including 6 head and neck squamous cell carcinoma (HNSCC)^20^, 8 multiple myeloma (MM), 2 oral squamous cell carcinomas (OSCC)^45^ and 4 ovarian cancer patients (OV)^46^ (**Table S5**).

We obtained SCCN profiles from the scRNA-seq data using the *inferCNV* program^47^ (**Methods**), which derives CNAs from averaged mRNA expression levels in consecutive 150-gene windows across the genome. We then obtained a MEDALT for each patient, including cells in both the primary and the metastasis samples. For comparison, we also performed transcriptomic trajectory analysis for each patient using Monocle v3.0^48^. Since the cells in the primary samples were most likely born before the cells in the metastasis (or relapse) samples, they should be arranged closer to the root of the lineage trees. Indeed, in the MEDALTs, the cells from the primary samples were placed significantly (one-side Wilcoxon rank-sum test, p=0.0098) closer to the root than the cells from the metastasis (or relapse) samples (**Fig. 4a**). In contrast, the pseudotime estimated by Monocle failed to significantly (one-side Wilcoxon rank-sum test, p=0.51) delineate the two types of cells (**Fig. 4b**). Meanwhile, cells in the MEDALT lineages had more homogenous SCCN profiles than those in the Monocle clusters (**Fig. 4c and Fig. S8, Methods**).

**Fig. 4.**
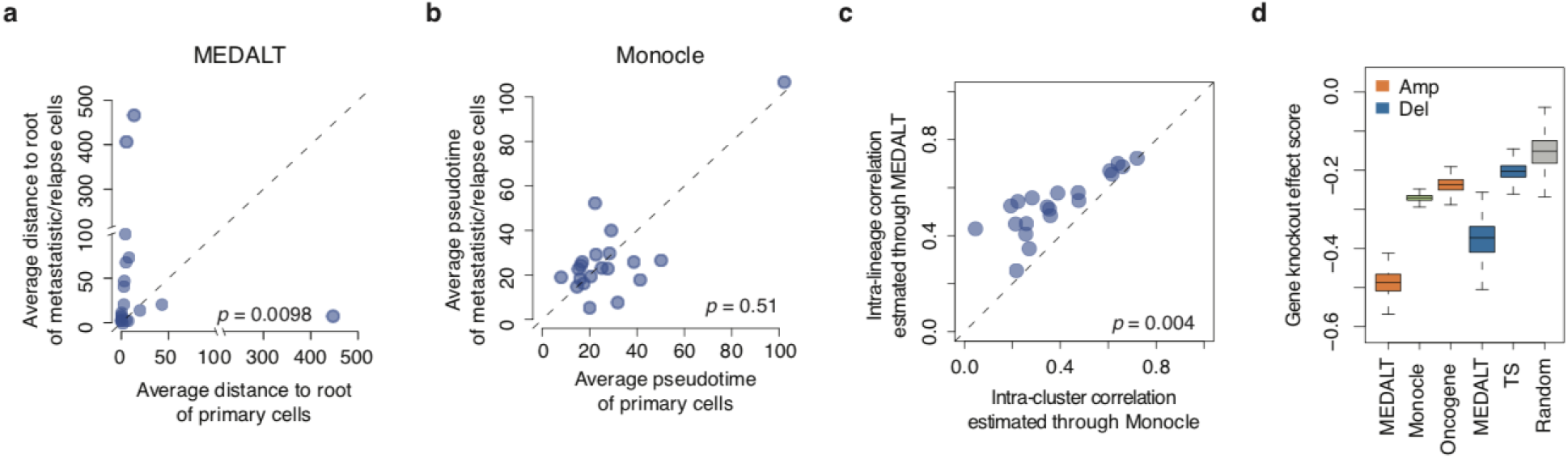
Application on scRNA-seq data from cancer patients. **a.** Average distance to root of the cells in the primary samples (X-axis) and those in the metastasis/relapse samples (Y-axis) estimated from the MEDALTs. **b.** Average pseudotime of the cells in the primary samples (X-axis) and those in the paired metastasis/relapse samples (Y-axis) estimated by Monocle. **c.** Pearson’s correlation coefficients between the SCCN profiles of the cells in the same lineages dissected from the MEDALTs (Y-axis) versus those in the same cell states clustered by Monocle (X-axis). Each dot in **a, b** and **c** represents a cancer patient. All the *p*-values were estimated by one-side Wilcoxon rank-sum test. **d.** Average DepMap gene-knockoff effect scores of the gene set identified by MEDALT and those by Monocle from the 20 patients. Also included as controls are 100 sets of 75 random genes, 234 oncogenes and 269 tumor suppressors (TS) in oncoKB and intOGen.

We performed cohort-level LSA on the MEDALTs and identified 75 fitness-associated genes (**Supplementary Data 4**, *p* value < 0.001), which included 45 amplified and 30 deleted genes from the 20 patients. In contrast, Monocle identified 3,412 differentially expressed genes between the cell clusters.

We found that the amplified genes identified by our approach are significantly more essential than those identified by Monocle (one-side Wilcoxon rank-sum test, **Fig. 4d**), based on the CRISPR screens of 524 cancer cell lines in the DepMap^37^.

## Discussions

Advances in single-cell technologies present new challenges and opportunities for making biological discovery. Single-cell studies often involve large numbers of cells, which are powerful at characterizing cellular heterogeneity, but small numbers of biological samples, which are underpowered for discovering common disease genes. It has been shown by recent genome-wide association analysis that it is possible to enable new discovery by performing association analysis at cell-type resolutions^49^. For cancer and genetic diseases driven by somatic mutations, being able to obtain genetic footprint at various time and conditions can enable discovery of genes responsible for disease progression and resistance to therapy.

However, it remains unclear what analytical strategies should be deployed to achieve the benefits. Even more challenging it gets when CNAs are being considered, as CNAs affect large regions of the genome and are difficult to trace using phylogenetics methods.

In our study, we demonstrated that it is possible to achieve the benefit by reconstructing copy number evolution history as a lineage tree, i.e., MEDALT, and performing permutation-based statistical analysis, i.e., LSA, to identify fitness-associated CNAs and genes.

We have learned several important lessons in our study.

First, it is important to perform accurate lineage tracing. Although the single-copy gain and loss model that we implemented in deriving MEDALTs is limited in complexity, it already performed substantially better than conventional phylogenetics algorithms such as MP that assumes infinite sites and NJ that employs naïve distance metrics, as shown in our simulation and in real data analysis. It is conceivable that further development of methodology that incorporates more complex genome evolution mechanisms such as chromothripsis^50^ can lead to better results.

An important goal was to represent convergent evolution that is likely prevalent in the lens of CNAs^10, 51^. Current phylogenetics algorithms strictly prohibit the expression of convergent evolution by disallowing an alteration to occur multiple times in a course of evolution^25^. As shown in our analysis of the TNBC patients, genes identified based on convergent evolution analysis (i.e., PLSA) had an even higher fraction of known cancer genes than those identified based on cohort-level single-lineage LSA. Our result suggests that examining convergent evolution is likely a key component towards fully unleashing the power of single-cell studies.

One important difference between MEDALTs and phylogenetics trees is that MEDALTs are minimal spanning trees that do not contain unobserved ancestral nodes. Representing evolution using minimal spanning trees instead of phylogenetics trees was our deliberate choice, as it allowed us to develop polynomial-runtime solutions that are scalable to real datasets containing thousands of cells. It also allowed us to conveniently implement biologically meaningful MED and enforce directionality constraints. Phylogenetics algorithms are likely effective when the numbers of cells are small and that the alterations are simple to trace. None of these conditions apply to available SCCN datasets that have CNAs evolving non-linearly in hundreds of cells. Moreover, we have shown in our simulation that for the purpose of detecting fitness-association alterations, our method outperformed phylogenetics approaches in a wide range of sample sizes.

A particular challenge in developing and evaluating computational lineage tracing methods, is the lack of exact ground truth. Although various experimental technologies have been developed^52, 53^, we are not unaware of any that can be applied to trace copy number evolution in patient samples. To circumvent this, we utilized *in silico* simulation that mimics several prevalent CNA mechanisms to evaluate the accuracies of the reconstructed lineages and fitness-associated alterations. We also utilized longitudinal datasets on which we knew the biological stages of the cells to evaluate the chronological accuracy of the inference results. Although these strategies are unlikely sufficient to validate all the edges and lengths in the trees, they are objective and sufficient to discriminate various approaches.

Second, it is important to control biases in statistical inference. It is challenging to detect fitness-associated genes, as CNAs often affect a large number of genes and that the sample sizes are often small. Passenger CNAs that occur naturally in non-functional regions such as those near fragile sites or repeats could easily cloud the discovery. In addition, lineage tracing algorithms are unlikely to be perfect and could introduce distinct biases. To address these challenges, we employed LSA, which randomly permutes SCCN profiles into different cells to reduce the noise introduced by background genomic variations. And we reconstructed trees from permuted datasets to alleviate biases introduced by the lineage tracing algorithms. These procedures appeared important to achieve the accuracy. Further exploration of different ways to permute the data and to estimate the background distribution will likely lead to better results.

We assessed the functional impact of the identified genes using cell-line CRISPR essentiality screen data. We confirmed that the set of fitness-associated, amplified genes discovered by our methods are significantly more essential than other control gene sets in cancer cell lines.

We also nominated novel genes that appear to have prognostic values in TCGA and the METABRIC datasets. These assessment strategies likely have false positives and negatives. Further comprehensive, well-controlled and targeted experiments will likely be required to fully assess the functional impact and clinical values of these genes.

Lastly, it was exciting to observe benefits of our methods on both the scDNA-seq and the scRNA-seq data. Although RNA-derived copy number profiles may not be as accurate as those derived from DNAs, previous studies^54^ suggested that they can reasonably distinguish tumor clones. Our study further revealed the value of scRNA-seq data in lineage tracing and supported the notion that genomic profiles, even approximations, are more accurate than transcriptomic profiles in determining biological timing of cells. Our results opened doors toward utilizing scRNA-seq as a platform to understand genetics underlying developmental processes and perform gene discovery.

## Methods

### 1. Inferring minimal event distance

We use a modified parsimony scoring method to score the distance between two copy number profiles, which can be considered as non-negative integer arrays. We assume a copy number alteration (CNA) (event) can affect adjacent genomic regions (one single entry or *k* adjacent entries in array) by increasing or decreasing their values by 1. We define the minimal event distance (MED) between two arrays *a* and *b* to be the minimal number of CNAs needed to transition from *a* to *b* (**Fig. S1a**).

We propose a greedy algorithm (**Table S1**) which guarantees to find an optimal solution within a runtime of *O*(*m*) (**Appendix**), where *m* is the size of the array^55^. We add an additional restriction that MED equals to infinity, if the copy number at any site is going from 0 to any other number.

### 2. Constructing minimal event distance aneuploidy lineage tree (MEDALT)

The optimal aneuploidy lineage tree is a rooted directed minimal spanning tree (RDMST) with the least number of CNAs. We use an implementation of Edmond’s algorithm to infer RDMST (**Table S2**). Our algorithm runs in *O*(*VE*), where *V* is the node set and *E* is edge set. That is approximately as *O*(*n*^3^), where n is the size of the node set.

### 3. Lineage speciation analysis

We propose a statistic routine named lineage speciation analysis (LSA), which performs permutation tests on the topology of MEDALT or phylogenetics trees to identify CNAs that are non-randomly associated with cellular lineage expansion in a developmental process. In LSA, we start from the root node and iteratively remove edges to obtain all possible lineages (subsets of cells). For the *i*-th lineage, we calculate a cumulative fold level (CFL) for the *j*-th CNA event that sums together the copy number alteration level in constituent cells (**Fig. S1c**).

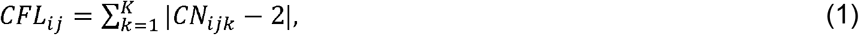

where *CN*_*ijk*_ is the copy number level in the *k*-th cell and *K* is the size of the lineage.

We treat the amplifications and deletions separately so that a region can be amplified in some samples but deleted in others. This is necessary because some oncogenes and tumor suppressors locate in close proximity and can get binned into the same regions.

We estimate the statistical significance of an observed CFL by comparing its value to a background distribution obtained through permutation (**Fig. S1d**). In the default mode, SCCN data are randomly shuffled by chromosomes into different cells (**Supplementary Note**). They are not further shuffled by sites within each chromosome, because chromosomal context plays an important role in determining where and how a CNA occur.

We construct a lineage tree for each permutated SCCN dataset and dissect each tree into a collection of lineages, from which we select the ones of identical (or very similar) size to the real lineage under test. We compute CFLs in these selected lineages using equation (1) and calculate an empirical p-value (tail probability) of the observed value:

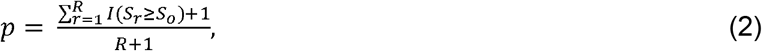

where *R* is the number of background lineages from the permutation data, *S*_*r*_, *S*_*o*_ are respectively the CFLs of the CNA event in the permutation and the real data.

### 4. Cohort-level LSA

In a cohort containing multiple individuals, we can estimate whether a CNA occurs non-randomly at the cohort (population) level. To do so, we construct meta-lineages by merging lineages dissected from different individuals and calculate a CFL for each meta-lineage through equation (1). We then estimate a statistical significance for each observed CFL through equation (2), based on a background distribution obtained from corresponding meta-lineages derived from individually permuted trees in the entire cohort (**Fig. S1e**).

### 5. Identifying parallel evolution event

The lineage speciation analysis (LSA) can be used to identify potential presence of parallel (aka. convergent) evolution (PLSA), i.e., finding CNAs that occur independently in multiple parallel lineages during the evolution of a cell population. We can assess the statistical significance of such events using the same permutation framework (**Fig. S1f**). Instead of examining each lineage independently, we deploy an algorithm that exhaustively searches for parallel lineages that are formed by disjoint sets of cells with identical CNAs or genes.

We then estimate the probability of observing such multi-lineage CNAs over random chance through permutation (as described above):

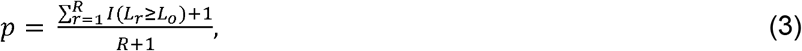

where *L*_*r*_, *L*_*o*_ are respectively the number of lineages containing the CNA of interest in the real and the permuted trees and *L*_*o*_ ≥ 2. *R* is the number of permutations. In this analysis, only CNAs tested positive in the LSA are being further considered for the PLSA.

### 6. Simulating single cell copy number evolution

#### Simulating cell birth-and-death process

In order to evaluate the accuracy of copy number lineage reconstruction, we implement a Markov process to simulate the cell growth under the influence of CNAs ^34, 56^. The simulation process starts from an ancestor cancer cell, which divides and dies at rate *b* and *d*, respectively. All the descent cells have the same division and death rates as do their ancestors, unless they are mutated.

The cell growth dynamics follow the following differential equation:

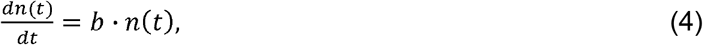

where *n*(*t*) is the number of cells at time *t*. We assume that there are one root and 2 children after the first division: *n*(0) = 1, *n*(1) = 2. That leads to *b* = 0.69 as the initial value based on equation (4).

The distribution of the time intervals Δ*t* between any two jumps in a Markov process with continuous time is exponentially distributed with the mean *E*(Δ*t*) = 1/(*b* + *d*)^57^. Here, we assumed *E*(Δ*t*) = 1 and the death rate *d* = 1 − *b*. When a jump occurs, it results in a birth with a probability *b*/(*b* + *d*) or a death with a probability *d*/(*b* + *d*). This cell birth-and-death process can be depicted as a rooted directed tree in which nodes are cells.

We simulated 100 independent runs, each of which has a population size of 200 cells.

#### Simulating the occurrence of CNA events

CNAs accumulate amongst tumor cells at an appreciable rate^58^. The CNAs in a cell at time t not only include the alterations it inherits from its parent, but also newly acquired ones from *t*_*i*−1_ to *t*_*i*_ (**Fig. S2a**). We assume that the CNA rate per site/region varies in several levels *μ* ∈ {0.02, 0.05, 0.1, 0.15, 0.2}^29^ and determine the number of CNAs (*K*) accumulating in Δ*t* based on a Poisson distribution (**Fig. S2a**):

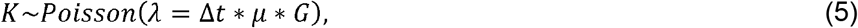

where *G* is the total number of sites/regions in the genome. In our simulation, we set *G* = 100.

#### Simulating genomic structural rearrangements

We assume that CNAs can be generated by various types of genomic structural rearrangements (GSR), such as terminal deletion (TER), interstitial deletion (DEL), unbalanced translocation (UT), tandem duplication (TD), inverted duplication (ID), and breakage fusion bridge (BFB)^27^. In addition, different GSRs could occur at differential rate in cancer^59, 60^. Thus, we determine the numbers of various GSRs based on a multinomial distribution^29^.

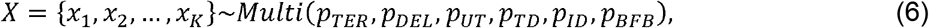

where we empirically set *p*_*TER*_ = *p*_*DEL*_ = 0.1, *p*_*UT*_ = 0.15, *p*_*TD*_ = 0.5, *p*_*ID*_ = 0.05, *p*_*BFB*_ = 0.1. We also required that 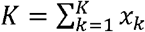 during the period of Δ*t* (**Fig. S2a**).

#### Simulating the location of a CNA

CNAs affect contiguous sites/regions in a chromosome. They often exhibit two modes: 1) focal, affecting a relatively small (<MB) region^61^ and 2) broad, encompassing large chromosomal regions (e.g., chromosomal arms)^62^. Broad CNAs often result from chromosomal mis-segregation during mitosis^58^, which is a hallmark of cancer. Both focal and broad CNAs are important in oncogenesis. While broad CNAs often manifest through dosage effects^13^, focal CNAs often target driver genes directly and result in protein structural changes^63^.

We determined the size *r* of a CNA in *X* by sampling a zero-truncated Geometric distribution:

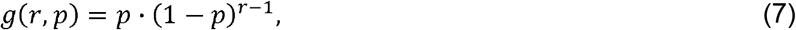

where *r* is the number of genomic sites/regions that a CNA occupies and *p* the probability that a region is affected by the CNA (**Fig. S2a**). We set *p* = 0.5 in our simulation.

We encode the simulated CNAs as sequences of non-negative integers in corresponding cells (**Fig. S2b**). Our model allows single-copy gains and losses. A copy number gain increases the corresponding values by 1 and a copy number loss decreases the values by 1 (**Fig. S2b**).

#### Simulating fitness-associated alterations

Some CNAs may themselves alter the fitness of a cell, or occur simultaneously with the driver mutations. We call them fitness-associated alterations (FAAs). We simulate the occurrence and the impact of FAAs in the evolution. At each generation, we determine if a FAA would occur through a Bernoulli distribution (*p* = 0.5). If a FAA occurs, we randomly select *τ* cells to carry the FAA, where *τ* follows a binomial distribution 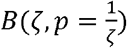 and *ζ* is the number of cells in the generation. The selected cells would increase their birth rates by *s*, which follows a uniform distribution *U*(0,1).

### 7. Constructing phylogenetic trees

We construct phylogenetic trees using the R package *phangorn^64^*, which implements widely used versions of the maximal parsimony, neighbor-joining and maximum likelihood approaches. To apply the maximal parsimony approach, the SCCN data are re-segmented by the collection of breakpoints detected in each cell, so that each column in the data matrix corresponds to a genomic interval that is uninterrupted by any GSR in any cell. The GSR breakpoints in individual cells are determined by the R package *copynumber* under default parameters. To apply the neighbor-joining approach, Hamming distances are calculated from each pair of the SCCN profiles. To applying the maximal likelihood approach, random trees are chosen as the initial solutions.

### 8. Estimating the accuracy of lineage partitioning

The cell birth-and-death process we simulate can be expressed as a rooted directed minimal spanning tree (RDMST). To compare RDMST with phylogenetic trees, we convert RDMSTs into dendrograms, which are fully comparable with the phylogenetic trees in that observed cells are represented as leaves in both types of representations^65^. From each dendrogram or phylogenetics tree, we calculate a metric, termed lineage partitioning accuracy (LPA), which measures how accurately cells are partitioned into lineages (subsets). Given a dendrogram, we performed lineage partitioning as follows:

We iteratively remove each branch in the dendrogram to obtain all the bi-partitions, i.e., the two disjoint subsets resulting from removing a branch. Each subset corresponds to a cellular lineage. All lineages can be described as a binary sequence *l* = {*c*_1_, *c*_2_, …, *c*_*N*_}, *c*_*i*_ = 1 if the *i*-th cell is in lineage *l* and *c*_*i*_ = 0, otherwise.

In the simulation experiments, the lineages partitioned from the simulated cell growth trees are considered as the ground truth. The LPA of a given MEDALT or phylogenetics tree is calculated as the fraction of lineages that exist in the ground truth over the total number of predicted lineages.

### 9. Accuracy of FAA detection in simulation

We randomly spike in FAAs in the simulation experiments, which are used as the ground-truth to assess the accuracy of the MEDALT and the phylogenetic trees. For each CNA, we calculate its *p* value through LSA and identify the minimal *p* value over all the lineages containing the CNA. We use −log (*minimal p*) as the prediction score. We then characterize the accuracy of each approach on FAA detection using AUC values, which are calculated by tallying the positive and the negative hits at various prediction score cutoffs from 0 to the maximal values.

### 10. Identifying significant CNAs using GISTIC

We apply the GISTIC algorithm on the simulated and the real SCCN datasets to identify significant CNAs^31^. The following steps are taken:

i. Calculate the occurrence frequency (*f*) and the amplitude (Δ) of each alteration;
ii. Define a G-score as a function of *f* and Δ: *G* = *f* × log_2_ (Δ + 2);
iii. Assess the statistical significance of each alteration by comparing the observed G-score to a background distribution of G-scores obtained from permuted (by regions) copy number profiles.

On the simulated datasets, we regard each cell as an individual sample and apply GISTIC at the cell level.

On the TNBC dataset, we average the SCCN profiles across the cells in each patient sample to create a pseudo-bulk copy number profile for each sample. We then run GISTIC on these pseudo-bulk profiles to identify significant CNAs, similarly to how GISTIC is applied in TCGA study.

### 11. Inferring copy number profiles from single cell DNA sequencing data

The SCCN profiles from single cell DNA-sequencing data are estimated using a variable binning method, as detailed in previous studies^18, 66^. Briefly, sequencing reads are counted in 11,927 genomic bins with variable start and stop coordinates, which are optimized to receive even read counts across the bins. The median genomic length spanned by the bins is 220 kbp. Cells with < 50 median reads per bin are excluded. Loess normalization is used to correct for GC bias^35^. Copy number profiles are segmented using circular binary segmentation (CBS)^67^ followed by MergeLevels^68^ to joint adjacent segments with non-significant differences in segment ratios (parameters alpha = 0.0001 and undo.prune = 0.05). Integer copy numbers are calculated by scaling segment ratios with average DNA ploidy determined by flow sorting indexes and rounding to closest integers ^18^.

### 12. Dissecting MEDALT into disjoint lineages

To characterize CNA rate variation and genetic organization of a cell population, we dissect it into disjoint lineages (cell subsets) based on the corresponding MEDALT. For each internal node *v* in MEDALT, the subtree rooted at *v* is denoted as *T_v_*, which consists of all the descendants of *v*. The number of nodes in *T*_*v*_ is denoted as *S*_*v*_, the size of the subtree. To ignore small lineages that cannot be confidently characterized, we set a minimal subtree size cutoff *s* (*s* = 5 in our analysis of the scDNA-seq and the scRNA-seq data) and define an internal node set *IV* = {*v*|*S*_*v*_ > *s*, *v* ∈ *V*}, where *V* represents the node set of the MEDALT. We arrange the node sets in *IV* in an increasing size order:

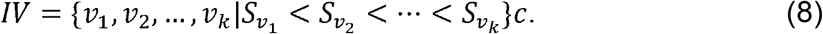

To obtain disjoint lineages, we remove the internal nodes that lead to redundant lineage assignments. For each *v*_*i*_ ∈ *IV*, 1 ≤ *i* ≤ *k*, its parent node *v*_*j*_(*j* > *i*) should exist in *IV*. If a parent node *v*_*j*_ has more than one child in *IV*, remove *v*_*j*_ from *IV*; otherwise, remove *v*_*i*_. We iterate through all the nodes in *IV* until no node can be removed. We then split the MEDALT into subtrees rooting at nodes remaining in *IV*. All the nodes that are not yet included are assigned into a control lineage.

### 13. Estimation of CNA rate and fraction of DDR loss

We estimate CNA rate in a lineage as the average number of CNAs, i.e., average MED, between the cells in the lineage. DNA damage repair (DDR) genes play key roles in maintaining genome stability. In our analysis, we download the list of DDR genes from Knijnenburg *et al*’s study^36^, based on which we estimate the proportion of DDR genes with copy number loss in each lineage.

### 14. Characterizing chromosomal level CNAs identified in TNBCs

In order to benchmark the accuracy of chromosomal (arm) level CNA detection in the TNBC data, we search biomedical literature exhaustively and create a list of chromosome-arm-level CNAs that have reported relevance to TNBC biology or clinical utilities (**Table S4**). We treat this list as the ground truth.

For each chromosomal level CNA in a lineage tree, we used the −log (*p*) estimated via the cohort LSA as its prediction score. We then estimate AUC values, respectively for the MEDALT, the MP and GISTIC approaches.

### 15. Inferring copy number profile from single cell RNA sequencing data

We use *inferCNV* (https://github.com/broadinstitute/infercnv) to identify somatic large scale chromosomal CNAs from single cell RNA sequencing (scRNA-seq) data^47^. *InferCNV* detects CNAs by exploring expression intensity of genes across positions of tumor genome in comparison to a set of reference “normal” cell. The CNAs at gene level relative to reference cell are estimated under default parameters. We calculate average relative CNA values in non-overlapping genomic bins, each consisting of 30 genes. Within each bin for each cell, we calculate an integer copy number by multiplying the relative CNA value by 2 (diploid) and then rounding the results off to closest integers.

### 16. Estimating genetic homogeneity

We compute a metric, called genetic homogeneity level (GHL) to compare the accuracies of MEDALTs with those of Monocle trajectories in tracing genetic evolution from scRNA-seq data. For each cell lineage (subset) partitioned from a MEDALT (section 12 Methods), we calculate pair-wise Pearson’s correlation coefficients between all the cells in the lineage, using gene-level copy number profiles inferred by *inferCNV*. We treat the mean correlation coefficient as the GHL of the lineage. Then average the GHLs across the lineages to obtain an overall GHL of the MEDALT.

Similarly, we calculate a GHL for a Monocle trajectory by averaging cluster-level GHLs estimated from cell clusters defined by the trajectory.

## Acknowledgements

This work was supported in part by the NIH [R01CA172652, U01CA217842], the CPRIT [RP180248], the MD Anderson Cancer Center Sheikh Khalifa Ben Zayed Al Nahyan Institute of Personalized Cancer Therapy grant [U54CA112970] and the NCI Cancer Center Support Grant [P30 CA016672]. This work was also supported by the Human Breast Cell Atlas Seed Network Grant (HCA3-0000000147) to N.N. and K.C. from the Chan Zuckerberg Initiative DAF, an advised fund of Silicon Valley Community Foundation.

