## Supplementary materials for "Single-cell copy number lineage tracing enabling gene discovery"

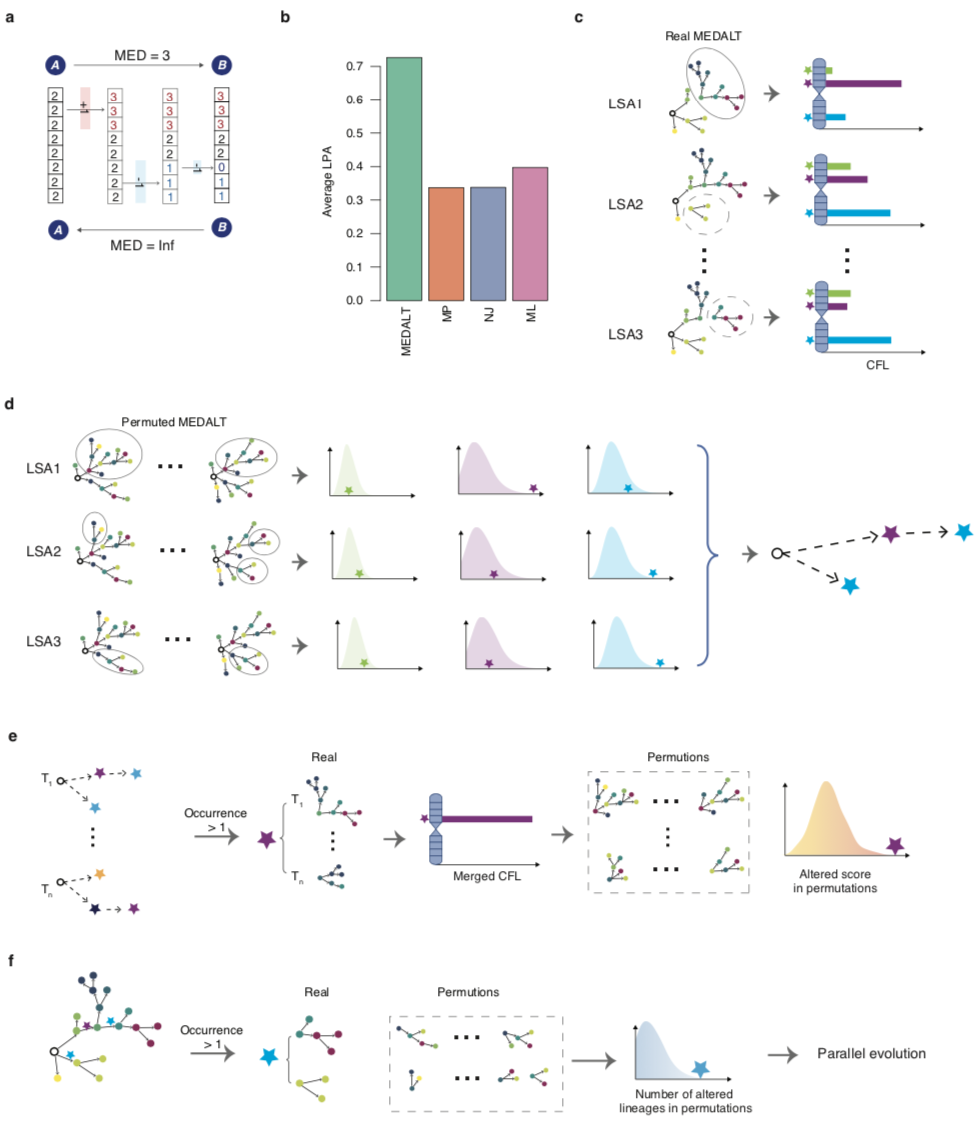

**Fig. S1** Methodology of the framework. **a.** Illustration of minimal event distance (MED) calculation. **b.** Average lineage partitioning accuracy (LPA) on 100 simulation datasets. **c**. Estimating lineage specific cumulative fold level (CFL). **d.** Estimating significance of CFL in an individual sample. **e.** Identification of non-random fitness-associated CNAs in a cohort of samples. **f.** Identification of parallel evolution CNAs in an individual sample.

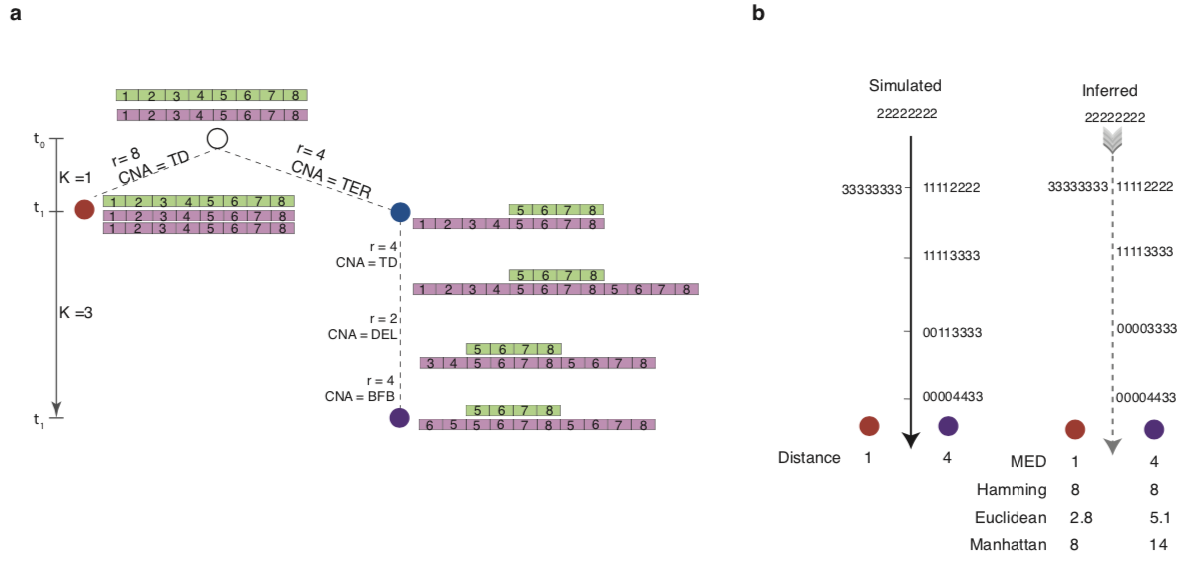

**Fig. S2**. Simulation of CNA evolution. **a.** Illustration of simulated genomic structural rearrangements in the evolution of a tumor. K represents the number of CNAs during $\Delta t$ period. r represents the number of adjacent regions which are affected by a CNA. TD: tandem duplication. TER: terminal deletion. DEL: interstitial deletion. BFB: breakage fusion bridge. **b**. Simulated and inferred copy number evolution distance between two genomes. Compared with MED are commonly used distance metrics Hamming, Euclidean and Manhattan.

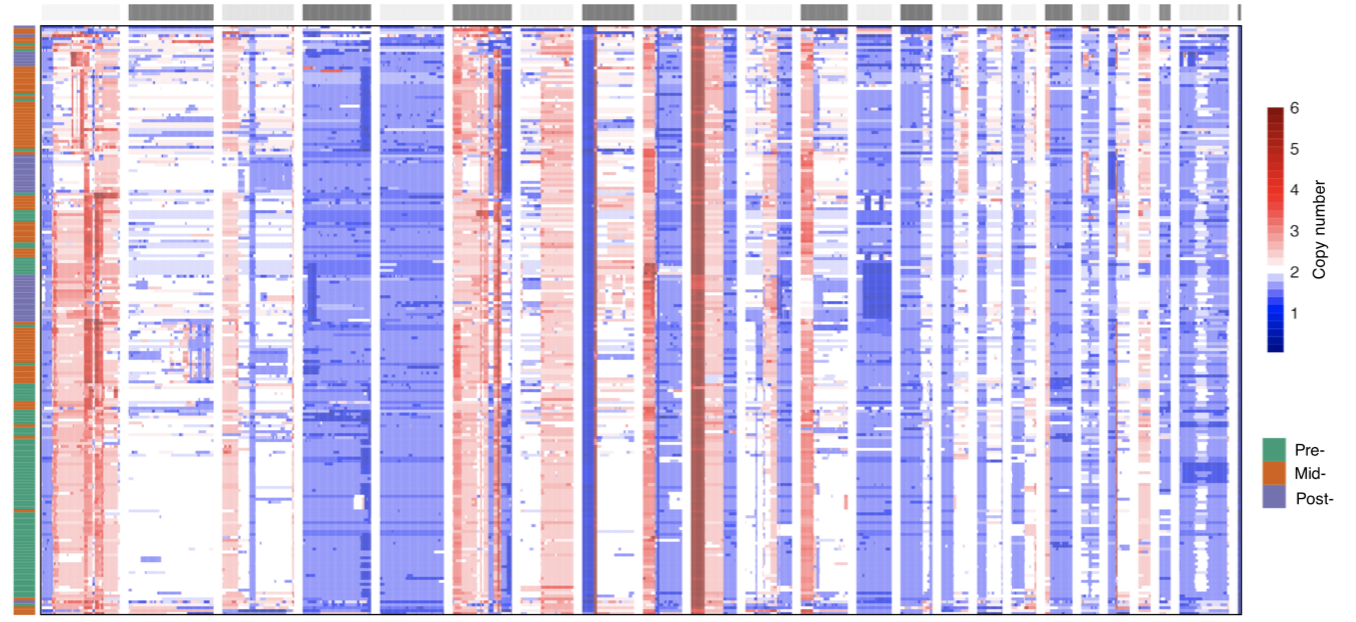

**Fig. S3**. SCCN profile of TNBC patient KTN102. Each row represents a cell from pre-, mid-, or post-treatment.

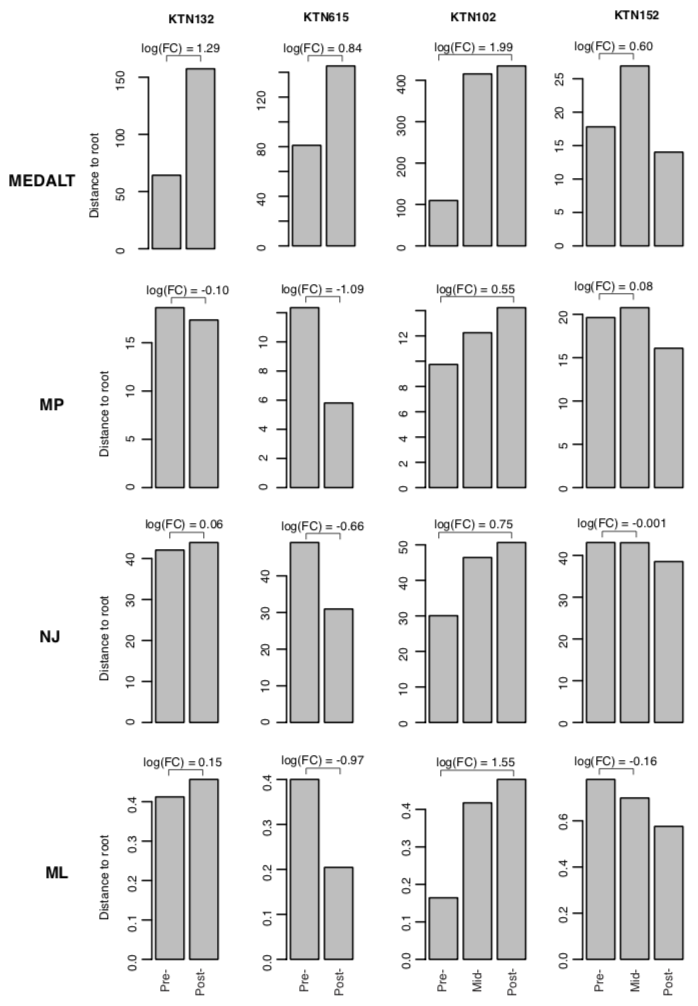

**Fig. S4**. Average distance between root node and cells from pre-, mid- or post-treatment based on MEDALT, maximal parsimony (MP), neighbor-joining (NJ) and maximum likelihood tree. FC refers to the fold changes between the average distance to root of the mid-/post- cells and that of the pre-treatment cells.

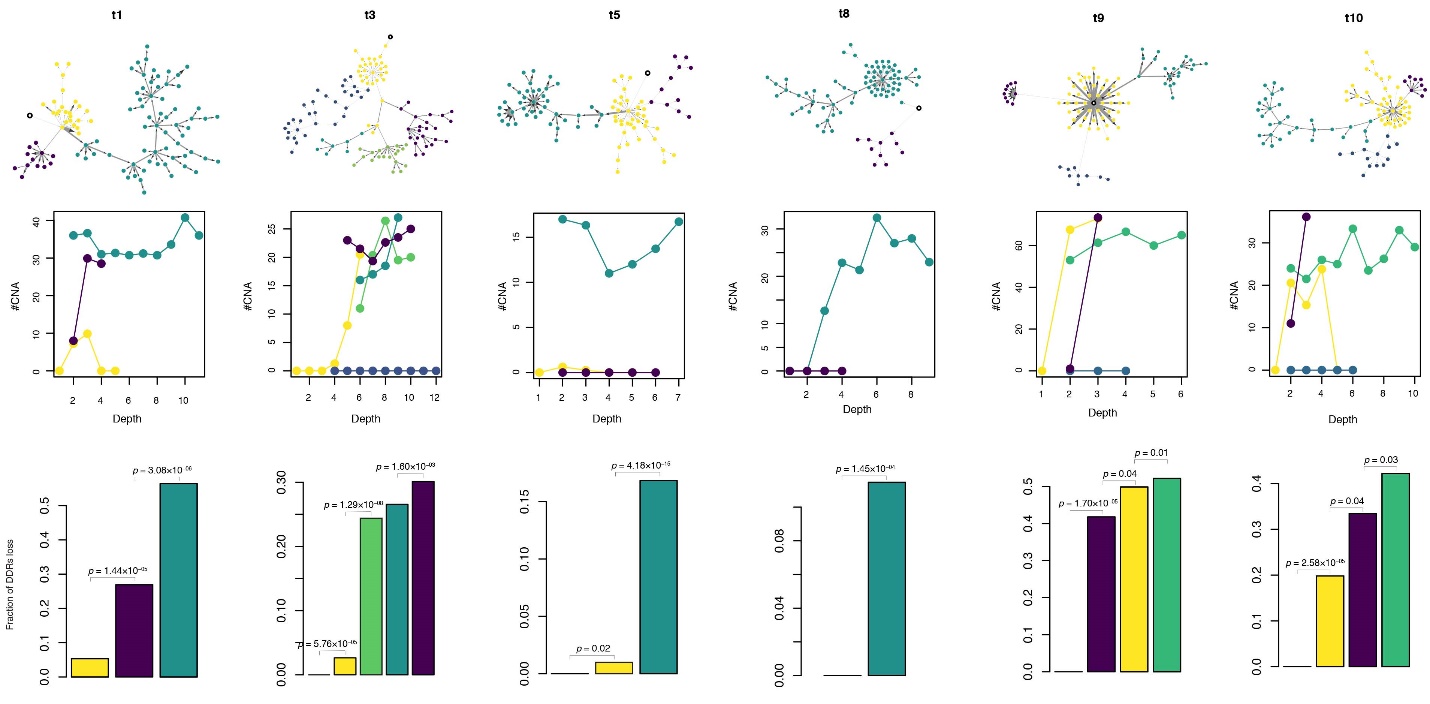

**Fig. S5**. Stratified average CNA rates and fractions of DDR genes loss among lineages (distinguished by colors) in 6 primary TNBC samples

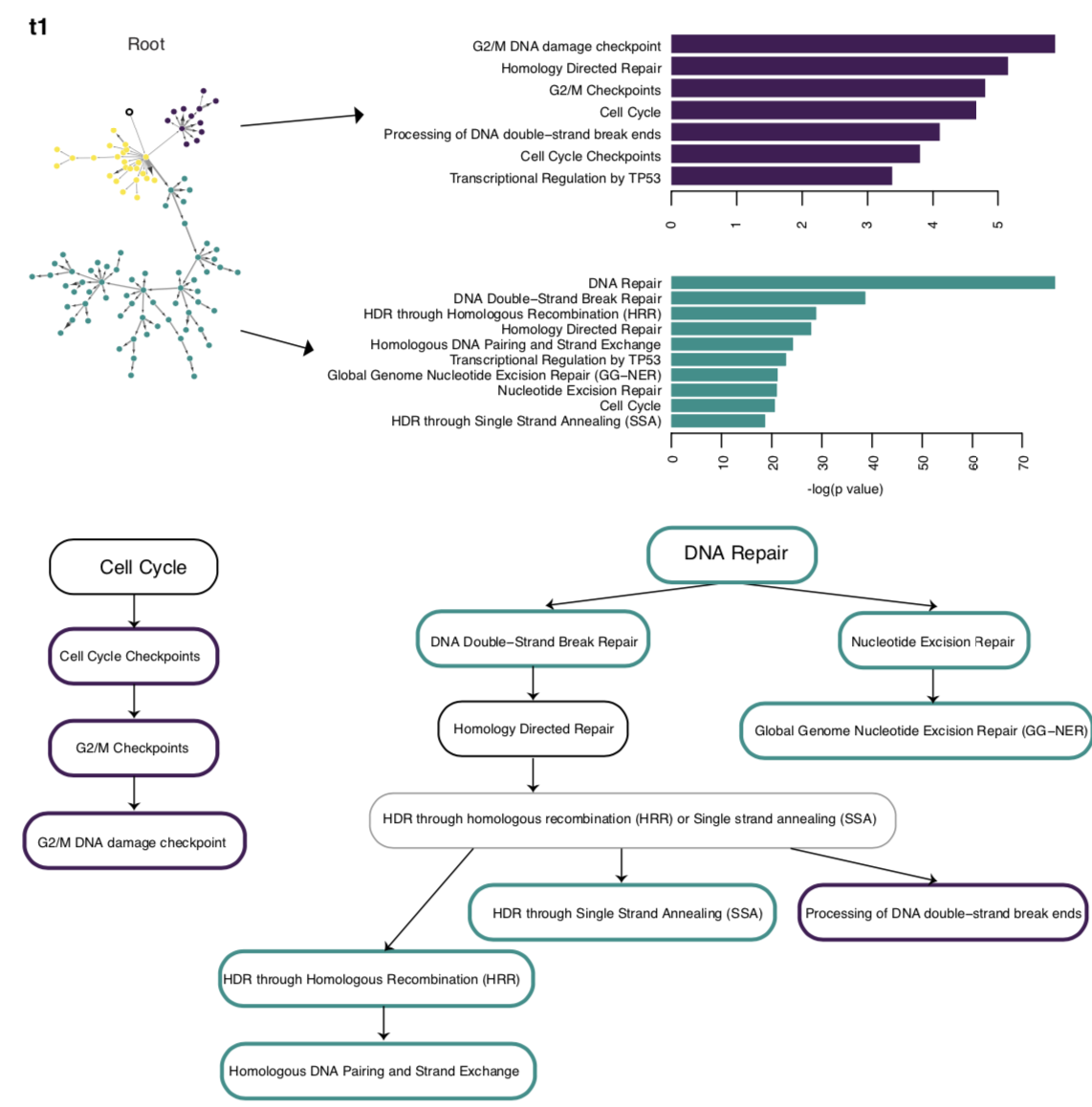
­

**Fig. S6**. Gene set enrichment analysis (GSEA) for genes identified by LSA in patient t1. Colors correspond to branches.

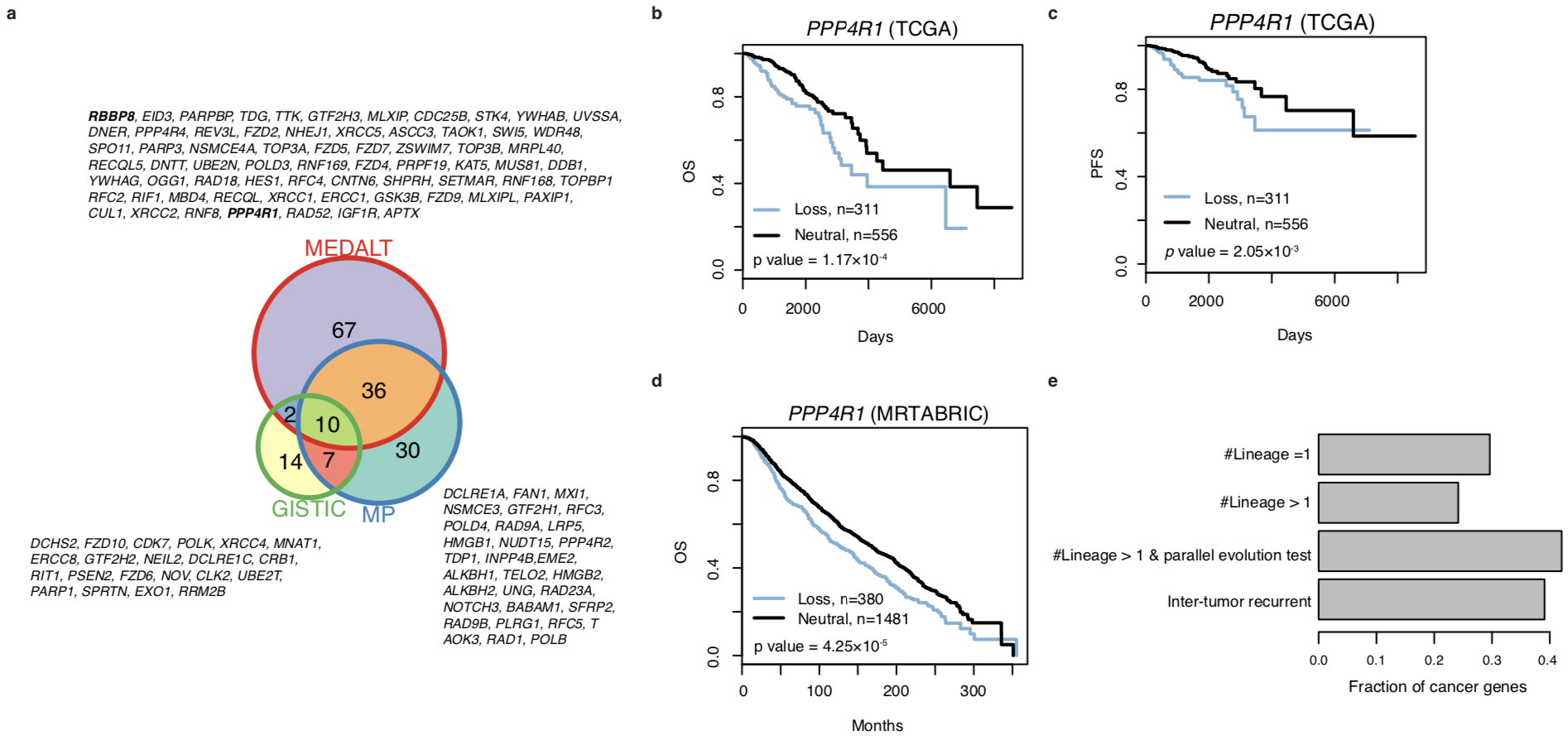

**Fig. S7.** Significant genes identified through cohort LSA from the TNBC scDNA-seq data. **a.** Venn diagram of the genes identified by the MEDALT, MP and GISTIC but not reported in oncoKB, COSMIC and intOGen. **b.** Overall survival (OS) analysis of breast cancer patients in TCGA. **c**. Progression free survival (PFS) analysis of breast cancer patients in TCGA. **d**. Overall survival analysis of breast cancer patients in the METABRIC. **e.** The fraction of cancer genes overlapping with events which were significant in single lineage (#Lineage = 1), multiple lineages (#Lineage > 1), parallel evolution test ((#Lineage > 1& PLSA < 0.001) and cohort LSA (inter-tumor recurrent).

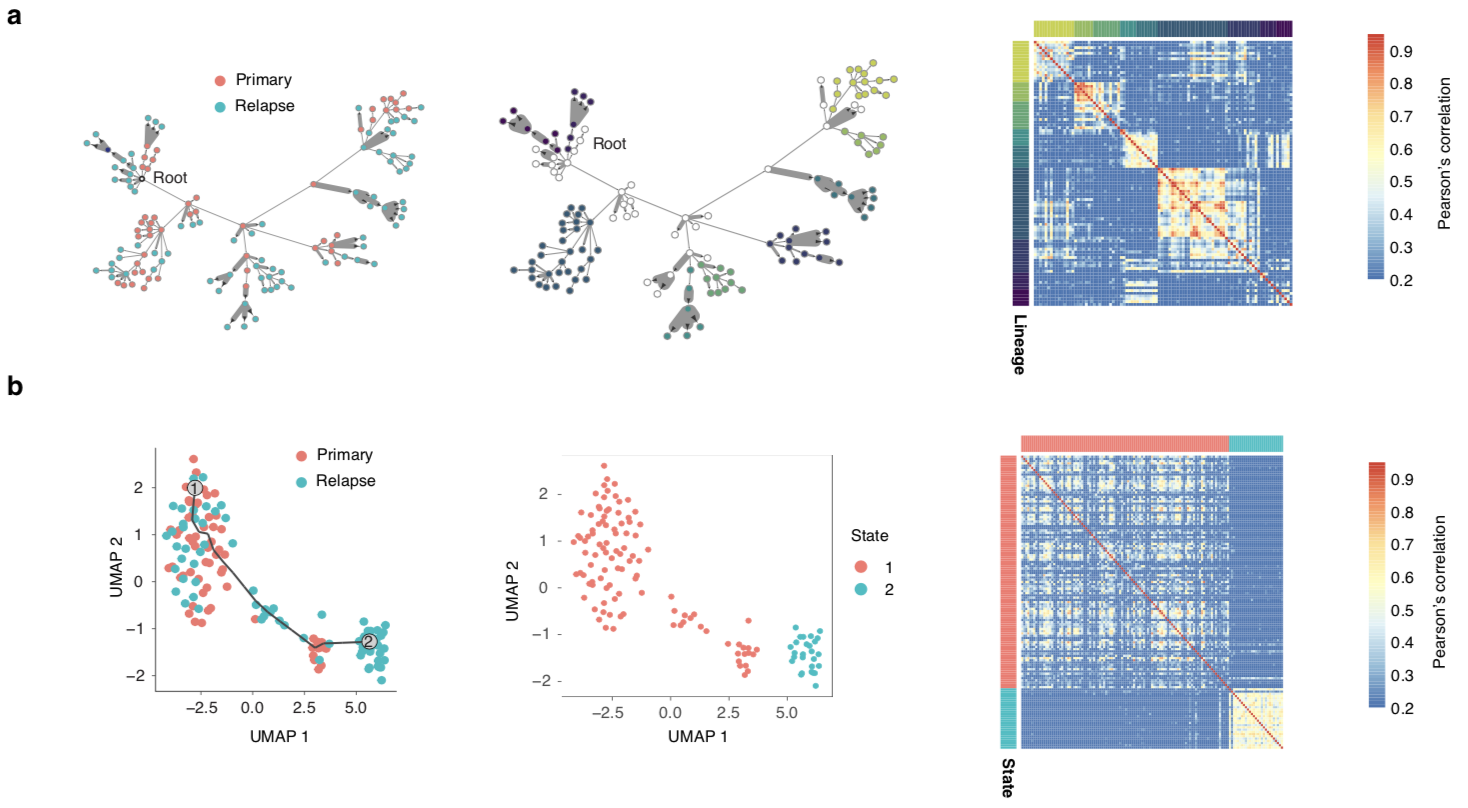

**Fig. S8.** Results of multiple myeloma patient 60359. **a.** Inferred MEDALT and heatmap based on Pearson’s correlation of the inferCNV profiles between cells ordered by lineages in MEDALT. **b.** Inferred trajectory from Monocle and heatmap of Pearson’s correlation of the inferCNV profiles between cells ordered by states defined by Monocle.

**Table S1. The algorithm for minimal event distance (MED) inference**

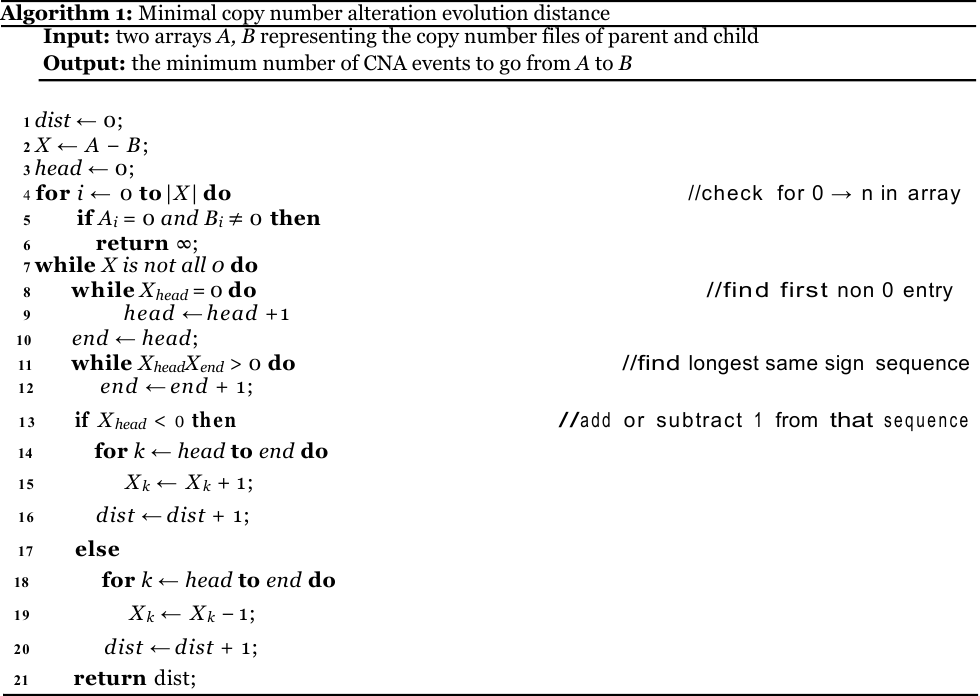

**Table S2. The algorithm for rooted directed minimal spanning tree reconstruction**

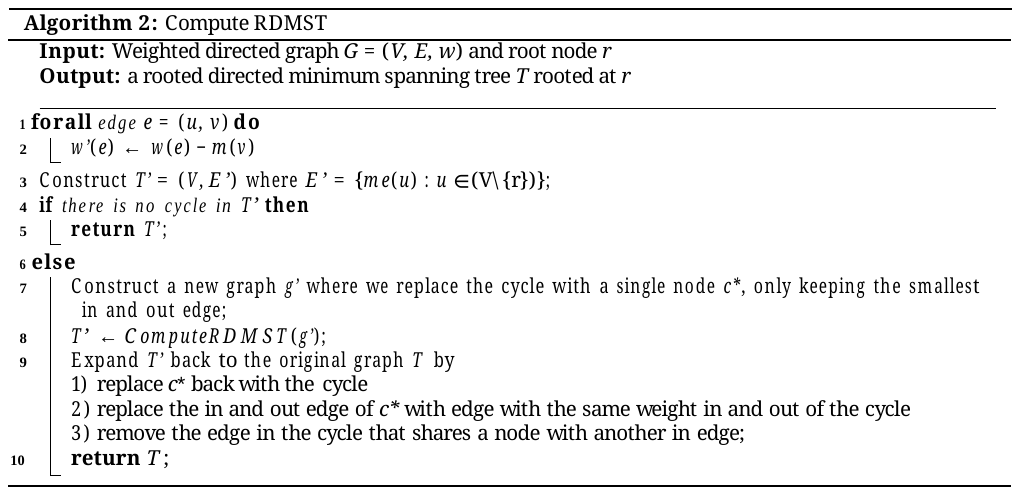

Where *m*(*u*) is minimum weight of all edge going into *u, me*(*u*) as the edge that has the minimum weight of all edges going into *u.*

**Table S3. Information on the TNBC data**

| **Patient ID** | **Sample size** | | |
| --- | --- | --- | --- |
|  | **Primary/Pre-treatment** | **Mid-treatment** | **Post-treatment** |
| t1 | 100 |  |  |
| t2 | 65 |  |  |
| t3 | 110 |  |  |
| t4 | 52 |  |  |
| t5 | 89 |  |  |
| t6 | 90 |  |  |
| t7 | 68 |  |  |
| t8 | 82 |  |  |
| t9 | 85 |  |  |
| t10 | 92 |  |  |
| t11 | 100 |  |  |
| t12 | 48 |  |  |
| KTN206 | 45 |  |  |
| KTN126 | 30 |  |  |
| KTN129 | 33 |  |  |
| KTN302 | 46 |  |  |
| KTN615 | 16 | 34 |  |
| KTN132 | 43 | 31 |  |
| KTN102 | 88 | 76 | 37 |
| KTN152 | 43 | 28 | 40 |

**Table S4. Annotation of the broad CNAs identified in TNBCs based on literature**

| **CNA** | **Region** | **Method** | **Cancer** | **Functional description** | **PMID** |
| --- | --- | --- | --- | --- | --- |
| Loss | Chr5q | MEDALT, MP | **Breast cancer**, prostate cancer, cervical cancer | Tumor growth and metastatic progression | 29562176, 29662167, 30022638 |
|  | Chr2q | MEDALT, MP | **Breast cancer** | More aggressive tumor type, moderately/poorly differentiated tumors. | 10861509, 8818658, 13680524 |
|  | Chr9q | MEDALT | **Breast cancer**, bladder cancer, head and neck cancer, lung cancer, liver cancer | Invasive tumor lesions, | 8550238, 11438741, 16149093, 8118798, 8306323, 10209942 |
|  | Chr3p | MP | **Breast cancer**, squamous cancer, ovarian cancer, Vulvar Carcinoma | reduces cell proliferation, poor survival | 29622463, 29558370, 30279231, 27170308, 10850424 |
|  | Chr17p | MEDALT | **Breast cancer** |  | [26328251](https://www.ncbi.nlm.nih.gov/pubmed/26328251), [25481507](https://www.ncbi.nlm.nih.gov/pubmed/25481507), 13680524 |
|  | Chr4p | MP | Meningiomas, metastatic thymic adenocarcinoma |  | 31591222, 28506304 |
|  | Chr21q | MEDALT, MP | **Breast cancer**, non-small cell lung cancer, oral cancer susceptibility | Associated with tumor suppressor | 13680524,12735585, 9743305, 9790505, 13680524, 9523199, 11259088 |
|  | Chr7p | MEDALT, MP | **Breast cancer** |  | 10861509 |
|  | Chr23p | MP | **Breast cancer** |  | 20101236 |
|  | Chr10q | MEDALT | **Breast cancer**, Brain tumor, head and neck cancer, oligodendrogliomas | Invasive phenotype, poor prognostic | 22429330, 14676808, 10404099, 11438486 |
|  | Chr16q | MEDALT, MP | **Breast cancer** | Invasive phenotype, tumor suppressor | 29558370, 19156836, 13680524, 14976537, 8971163, 16280054, 21706489, 23348384 |
|  | Chr22q | MEDALT, MP | Breast cancer, colorectal cancer, glioblasma, gastrointestinal stromal tumor, insulinoma | Tumor suppressor | 10850424, 13680524, 29665859, 14712485, 10861509, 15580284, 11739439 |
|  | Chr8p | MEDALT, MP | **Breast cancer**, Prostate cancer | Poor prognosis | 27611943, 13680524 |
|  | Chr15q | MEDALT, MP | **Breast cancer**, bladder cancer | More aggressive tumor type, metastatic carcinoma | 8649814, 14633686 |
|  | Chr4q | MEDALT | **Breast cancer**, colon cancer, multiple myeloma | Worse outcome | 9743305, 21717218, 10446964, 17077331 |
|  | Chr12q | MEDALT | **Breast cancer**, pancreatic cancer |  | 19602461, 15300227 |
|  | Chr20p | MEDALT | Colorectal cancer |  | 12490103, 19359472 |
|  | Chr6q | MEDALT, MP | **Breast cancer** | Tumor suppressor | 9155053, 10778973 |
|  | Chr17q | MEDALT | **Breast cancer,** non-small cell lung cancer |  | 7671234 |
|  | Chr5p | MEDALT | **Breast cancer** |  | 16570289 |
|  | Chr18q | MEDALT, MP | **Breast cancer** |  | 13680524 |
| Gain | Chr8q | MEDALT, GISTIC, MP | **Breast cancer**, melanoma, kidney cancer | More aggressive tumor phenotype, high risk of distant metastases and greater tumor size. | 10861509, 11514955, 22605478, 31645765 |
|  | Chr6p | MEDALT, MP | **Breast cancer**, melanoma, liver cancer, ovarian cancer, glioblastoma | More aggressive tumor phenotype,  advanced or metastatic disease,  poor prognosis. | 10861509, 11514955, 16790693 |
|  | Chr3q | MEDALT, MP | **Breast cancer**, lung cancer | More aggressive tumor phenotype. | 10861509, 18317062 |
|  | Chr7q | MEDALT, MP | **Breast cancer**, colorectal cancer | metastasis | 29558370, 26894854 |
|  | Chr10p | MEDALT, MP | colon cancer |  | 29580161 |
|  | Chr20q | MP | Colorectal cancer, pancreatic cancer |  | 22860045, 29169336, 6568296, 28991255, 29991641, 26894854 |
|  | Chr9p | MEDALT, MP | Multiple cancer |  | 29212506 |
|  | Chr12p | MEDALT, MP | **Breast cancer**, pancreatic cancer | Docetaxel resistance, Carboplatin Sensitivity, aggressive phenotype | 31213465, 30873387 |
|  | Chr5p | MP | Cervical cancer, head and neck cancer, renal cell carcinoma |  | 18559093, 31427592, 19521957 |
|  | Chr6q | MEDALT | **Breast cancer** |  | 17925008, 24969692 |
|  | Chr16p | MP |  |  |  |
|  | Chr1q | GISTIC | **Breast cancer** |  | 14976537, 17060936 |
|  | Chr10q | MP |  |  |  |
|  | Chr12q | MP | Classical lobular carcinomas |  | 18473330 |
|  | Chr14q | MEDALT, MP |  |  |  |
|  | Chr18p | MEDALT, MP |  |  |  |
|  | Chr2q | MP |  |  |  |
|  | Chr20p | MP | **Breast cancer** |  | 12755492 |
|  | Chr4q | MP |  |  |  |
|  | Chr11q | MEDALT, MP |  |  |  |
|  | Chr16q | MEDALT |  |  |  |

**Table S5. Information on the scRNA-seq data**

| **Cancer** | **Patient ID** | **Sample size** | |
| --- | --- | --- | --- |
|  |  | **Primary** | **Metastasis/Relapse** |
| HNSCC | HN5 | 112 | 20 |
|  | HN6 | 80 | 44 |
|  | HN20 | 572 | 90 |
|  | HN25 | 61 | 148 |
|  | HN26 | 127 | 140 |
|  | HN28 | 70 | 68 |
| OV | HG1 | 227 | 250 |
|  | HG2F | 253 | 165 |
|  | HG3 | 25 | 6 |
|  | LG2 | 34 | 24 |
| OSCC | HN120 | 278 | 270 |
|  | HN137 | 413 | 155 |
| MM | 60359 | 51 | 82 |
|  | 27522 | 915 | 85 |
|  | 47491 | 640 | 596 |
|  | 56203 | 931 | 69 |
|  | 57075 | 177 | 823 |
|  | 58408 | 112 | 225 |
|  | 59114 | 33 | 323 |
|  | 81012 | 421 | 512 |

**Supplementary Note Establishing statistical significance in LSA**

We proposed lineage speciation analysis (LSA) to delineate fitness-associated alterations (FAAs) from those passenger alterations. Passenger CNAs that occur naturally in non-functional regions such as those near fragile sites or repeats could easily cloud the discovery. To reduce biases of FAA identification, we randomly permuted SCCN profiles into different cells to reduce the noise introduced by background genomic mutations. And we reconstructed trees from permuted datasets to alleviate biases introduced by the tree inference algorithms. For each CNA in each candidate lineage, we assessed the significance of observed cumulative fold level (CFL) based on background distribution established from lineages of similar sizes in the permuted trees. To evaluate the performance of LSA for controlling biases in statistical inference, we explored two additional ways to estimate the background CFL distributions:

Control 1: Rather than reconstructing a tree from each permuted SCCN matrix, estimate CFLs using the by-chromosome-permuted SCCN matrix and the tree reconstructed from the real data.

Control 2: Same as 1 except using the SCCN matrix permuted by chromosomal bins within each cell (similar to GISTIC) instead of by chromosomes across different cells.

On the 100 simulated datasets with spiked-in FAAs, we assessed statistical significance of the observed CFL with respect to the background distributions established from the two alternative ways. We calculate an empirical p-value for an observed by tail probability:

$p= \frac{\sum_{r=1}^{R} I(S_{r}\geq S_{o})+1}{R+1}$,

where $R$ is the number of permutations. We generated 1000 permuted datasets. $S_{r}, S_{o}$ are respectively the CFL of the CNAs calculated in the real and the permuted SCCN profiles.

For each CNA, we calculated the minimal *p* value over all the lineages containing the given CNA, and used $-log(minimal p)$ as its prediction score. We estimated AUC value based on the prediction score to evaluate the performance of FAA identification.

We found that the MEDALT approach using permuted tree as background achieved substantially better detection performance than both of the alternative ways of performing permutation **(Fig. S9)**. Control 1 was more effective at reducing the background noise than Control 2. Reconstructing lineage tree on each permuted dataset, as we described in Methods, further alleviates biases in statistical inference. The benefits appeared robust w.r.t. down-sampling until the number of cells dropped below 60 (**Fig. S9**).

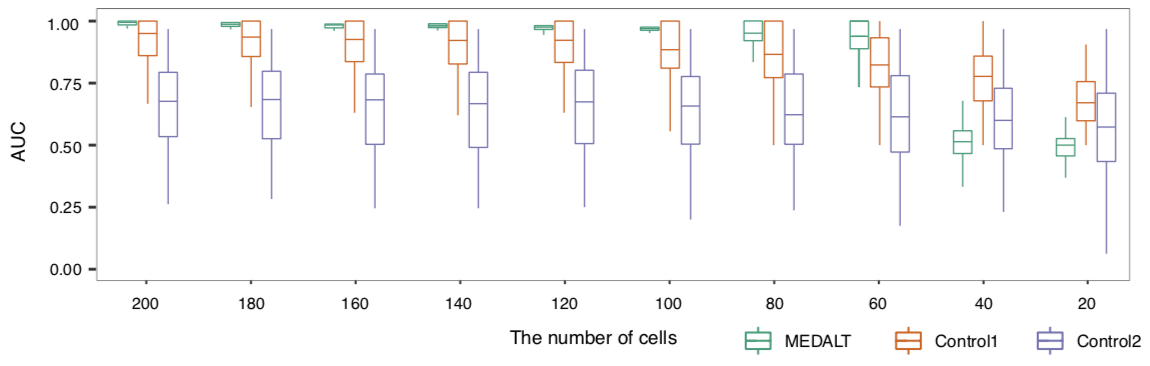

**Fig. S9.** AUCs of FAA identification based on 100 synthetic datasets.

**Appendix**

*Algorithm of minimal evolution distance (MED) inference*

Here we provide a theoretical proof and runtime analysis for the algorithm purposed in Table S1

$X=[x_{1},x_{2},\ldots x_{m}]$ and $Y=[y_{1},y_{2},\ldots y_{m}]$ are integer arrays of length $m$. The entries $x_{i}$ and $y_{i}$ indicate the number of DNA copies of position $i$ in cell $X$ and $Y$. The formulations here only refer to one chromosome but clearly generalizable to settings of multiple chromosomes.

An operation, or event, acting on array $X$ increases (amplification) or decreases (deletion) values by 1 in a contiguous segment of $X$. The minimal evolution distance (MED) between $X$ and $Y$ is the minimal number of events. We add an additional restriction that once a position $l$ has been lost, i.e. $x_{l}=0$, it can never be regained or further deleted. Thus, MED equals to infinity if the copy number at any site is going from 0 to any other number.

The following two lemmas are introduced based on the problem definition.

Lemma 1. For all positions $1\leq i\leq m$, different events may affect the same position, but opposite events cannot be applied on the same position.

Lemma 2. Given a sequence of events $C=\left( c_{1},c_{2},\ldots,c_{t} \right)$ that yields $Y$ from $X$, events in the sequence are commutative.

Let $C$ be an optimal copy number transformation (CNT) from $X$ to $Y$, our algorithm can find an equivalent sequence of events $C'$ which has the same length as $C$.

**Proof.** Given Lemma 2, we sort $C$ both by coordinates and decreasing length of segments affected by the event. $C=\left( c_{1},c_{2},\ldots,c_{t} \right)$, in which $c_{j}=(s_{j},h_{j},w_{j})$ indicate the event, $s_{j}$ and $h_{j}$ are start and end positions of the segment affected by the corresponding event, $w_{j}$ is either 1 for amplification or -1 for deletion.

$\forall j (1\leq j\leq t)$,

1) if $h_{j}<s_{j+1}$, the segments corresponding to $c_{j}$ and $c_{j+1}$ are disjoint. Thus, we can simply define ${c'}_{j}=c_{j},{c'}_{j+1}={c'}_{j+1}$.
2) if ${s_{j}\leq s_{j+1}\leq h}_{j}\leq h_{j+1}$, the segments corresponding to $c_{j}$ and $c_{j+1}$ are overlapped. If $w_{j}= w_{j+1},$ we define ${c'}_{j}=\left( s_{j},h_{j},w_{j} \right), {c'}_{j+1}=(h_{j}+1,h_{j+1},w_{j+1})$. If $w_{j}\neq w_{j+1}$, given Lemma 1, we define ${c'}_{j}=\left( s_{j},s_{j+1},w_{j} \right), {c'}_{j+1}=(h_{j},h_{j+1},w_{j+1})$.
3) if ${s_{j}\leq s_{j+1}\leq h}_{j+1}\leq h_{j}$, the segment corresponding to $c_{j+1}$ is included in the one corresponding to $c_{j}$. If $w_{j}= w_{j+1},$ we define ${c'}_{j}=c_{j},{c'}_{j+1}={c'}_{j+1}$. If $w_{j}\neq w_{j+1}$, given Lemma 2, we define ${c'}_{j}=\left( s_{j},s_{j+1},w_{j} \right), {c'}_{j+1}=(h_{j+1},h_{j},w_{j})$.

Finally, we obtain the solution $C'$ which yields $Y$ from $X$ via the same number of events as optimal CNT.

We’ve shown that we can transform all sequence of events into an equivalent sequence where the segments are sorted by start position and then by length. Then we want to show that our algorithm gives a solution that is as good as any optimal solution, hence giving the correct length.

Since no position is increased and decreased, the total amount of change needed is $|X-Y|$. Since we sort the events by their start position, then consider the leftmost location where there is still change to be made ($X_{i}\neq Y_{i}$), by our sorting rule, the next event will affect this position. Since our algorithm always looks for the longest subsequence of change to make, we will make more or equal change than the optimal solution. So the event length of our solution will be less equal to the optimal solution and that our solution is also optimal.

Runtime Analysis

The algorithm (Table S1) solves optimal CNT in time $O(m)$, where $m$ is the length of input array.

Input two arrays $X=[x_{1},x_{2},\ldots x_{m}]$ and $Y=[y_{1},y_{2},\ldots y_{m}]$, we assume the CNT is from $X$ to $Y$.

We set initial $MED = 0$.

If $\exists i \left( 1\leq i\leq m \right)$ such that $x_{i}=0$ and $y_{i}\neq0$, $MED= \infty$. This is performed in linear time.

Otherwise, we calculate the distance $D=Y-X=[d_{1},d_{2},\ldots,d_{m}]$. This is performed in constant time $O(1)$.

For each position $i$, $\left| d_{i} \right|$ events are needed from $X$ to $Y$. If $d = 0$, we skip the corresponding positions. We find a sequence of events $C=\left( c_{1},c_{2},\ldots,c_{t} \right)$, $c_{j}=(s_{j},h_{j},w_{j})$ such that {$d_{s_{j}},\ldots,d_{h_{j}}\}$ corresponding to the longest contiguous entries having the same change direction. $w_{j}=max\{{|d}_{s_{j}}|,\ldots,|d_{h_{j}}|\}$. Thus, $MED = \sum_{j=1}^{t} w_{j}$. The MED is calculated in linear time $O(m)$.

Therefore, the algorithm runs in linear time $O(m)$.
